# FACT-(H3-H4) complex stimulates Pol α activity to coordinate DNA synthesis with nucleosome assembly

**DOI:** 10.1101/2024.08.08.607175

**Authors:** Wenshuo Zhang, Jiawei Xu, Jiayi Yang, Guojun Shi, Jiale Wu, Ning Gao, Jianxun Feng, Qing Li

## Abstract

Deficiencies in replication-coupled (RC) nucleosome assembly often lead to reduced DNA replication rate, but the precise mechanism underlying this process remains unsolved. Here, we discovered that H3-H4, but not H2A-H2B, mediates the interaction between FACT and the primase-polymerase complex DNA Pol α. This interaction stimulates the DNA polymerase activity of Pol α, and is indispensable for Okazaki fragment synthesis and replication fork progression. Moreover, the Pol1-N domain of Pol α provides a specific binding site for FACT and H3-H4. Furthermore, CAF-1 and Rtt106-mediated replication-coupled nucleosome assembly pathways regulate this interaction. Together, we propose that the FACT-(H3-H4)-Pol α interaction acts as a “Pre-Warning System” that regulates DNA replication, ensuring proper coordination between DNA synthesis and nucleosome assembly.

## Main Text

DNA replication is one of the most sophisticated cellular processes, requiring precise coordination at the replication fork (*1, 2*). The replisome, a multi-protein complex, unzips the parental DNA helix and duplicates the separated strands during DNA replication (*3, 4*). The physical interactions between replisome components ensure a coordinated framework, facilitating efficient DNA synthesis (*5*). In eukaryotes, the replisome operates within a chromatin environment, adding an additional layer of complexity (*6–9*). Limiting the time that DNA remains naked is crucial to avoid nuclease attacks, which can lead to genome instability---a hallmark of cancer (*6, 8, 10*). To prevent the exposure of naked DNA, a surveillance mechanism should be in place to monitor the synchronization between nucleosome assembly and DNA replication speed, thereby avoiding severe issues caused by any mismatch between these two processes.

Physical interactions between replisome components and chromatin factors provide a solid foundation to orchestrate activities at the replication fork within the chromatin context. Accumulated evidence has shown that some replisome components can directly bind to histones and/or histone chaperones (*8, 11*). While these interactions facilitate replication-coupled (RC) nucleosome assembly by regulating new histone deposition or parental histone transfer, the precise mechanism by which these interactions fine-tune DNA synthesis remains unsolved. This is especially relevant given that a shortage of histone synthesis or the absence of certain RC histone chaperones like CAF-1 (<u>c</u>hromatin <u>a</u>ssembly <u>f</u>actor 1), Rtt106 (regulator of <u>T</u>y1 transposition <u>p</u>rotein 106 and FACT (<u>fa</u>cilitates <u>c</u>hromatin transaction) often results in a reduced replication rate (*12–20*).

In particular, multiple replisome components can interact with FACT and often form a module including replisome component(s)-histones-FACT(*21–23*). Among the replisome components involved in interacting with the FACT-histones complex, some are enzymes, such as the replicative helicase Mcm2-7 and the primase-polymerase complex DNA polymerase α (Pol α) (*23–25*). FACT is essential for helicase activity on a nucleosomal template but not on naked DNA in cell-free systems (*26, 27*), indicating that FACT assists helicase activity by promoting the disassembly of parental nucleosomes and recycling parental histones. Pol α is responsible for synthesizing the initial RNA-DNA hybrid primer required for DNA replication. Eukaryotic Pol α is typically composed of four subunits: the largest catalytic polymerase subunit Pol1 (POLA1 in humans), a regulatory B subunit Pol12 (POLA2 in humans), and two primase subunits Pri1 and Pri2 (PRIM1 and PRIM2 in humans) (*28, 29*). Primase synthesizes 8-10 nucleotides (nt) of RNA that are then transferred to Pol1 for a limited extension to a total primer length of about 20-35 nt (*30–35*). FACT interacts directly with Pol1 (*36, 37*), and Pol1 also possesses a histone-binding ability and binds both H3-H4 and H2A-H2B (*38–41*). However, whether and how the interactions of Pol α with FACT and/or histones might affect Pol α’s activities remain to be determined.

## Results

### Histone H3-H4, but not H2A-H2B, promotes the interaction between FACT and Pol α

To test whether FACT and histones can affect Pol α activity, we first characterize the interaction among FACT, histones, and Pol α based on fluorescent assays in living cells (Fig. 1A and fig. S1A). Since FACT directly interacts with Pol1, the largest subunit in Pol α (*36, 37*), we focused on Pol1 in this study. Using a bimolecular fluorescence complementation (BiFC) assay, the fluorescence signal arises from the complementation of two complement non-fluorescent mVenus fragments (e.g., Vn173 and Vc155) brought together by their respective fusion partners. In yeast, FACT consists of two essential subunits, Spt16 and Pob3. We tagged the Spt16 subunit with the N-terminus of mVenus (Spt16-Vn173), while the Pol1 with the C-terminus of mVenus (Pol-Vc155). The BiFC signals indicate that Spt16 interacts with Pol1 (fig. S1B and C, Spt16::Pol1), and both Spt16 and Pol1 interact with histone H3-H4 (fig. S1, B, D and E).

**Fig. 1.**
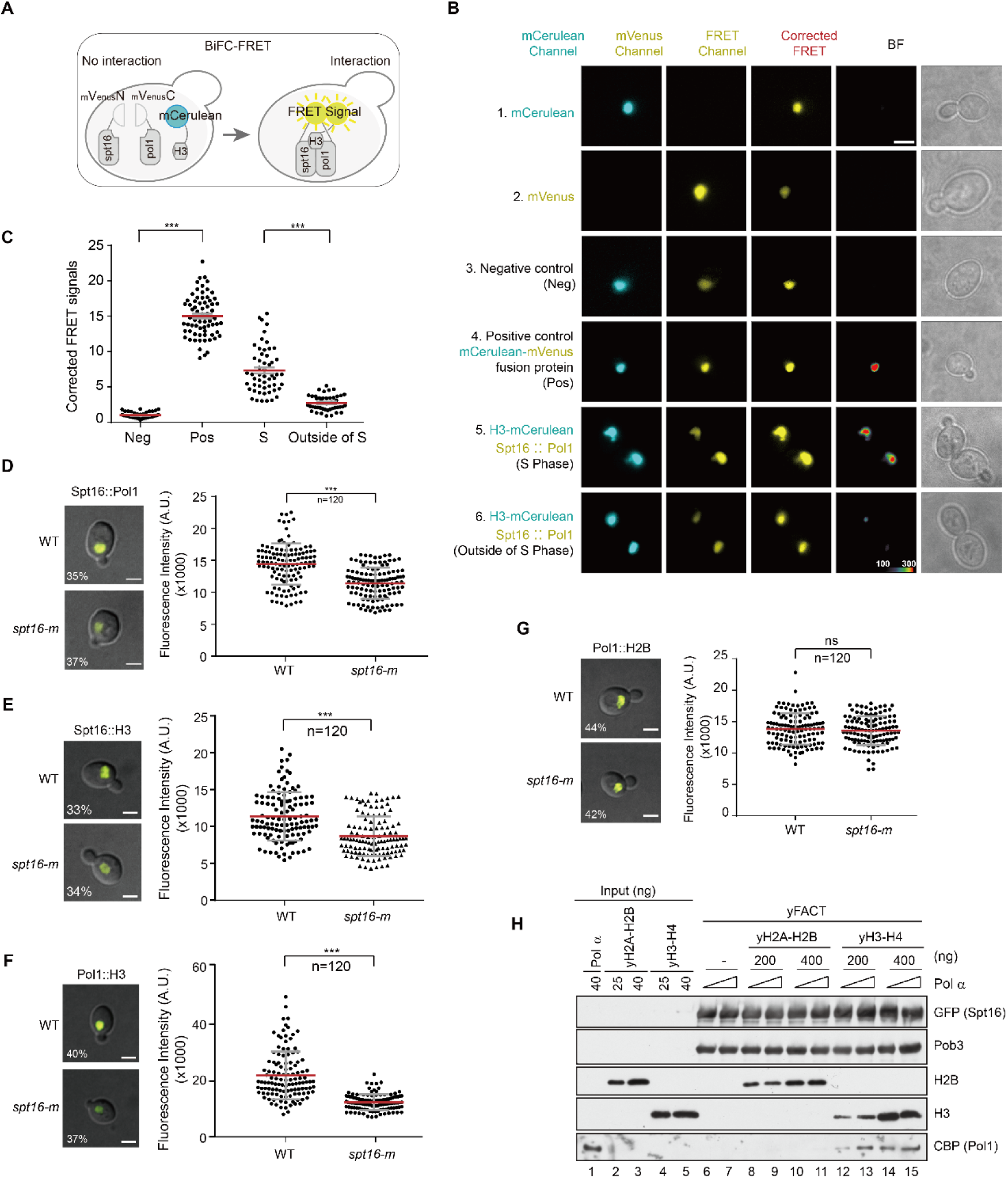
H3-H4, but not H2A-H2B, promotes the interaction of FACT with DNA polymerase α. (**A**) Schematic of BiFC-FRET method. N- and C-terminal of mVenus were fused to Spt16 and Pol1, respectively. Vn173 and Vc155 would reconstruct a functional mVenus when two fusion proteins interact with each other. When H3-mCerulean and the reconstructed FACT-Pol1-mVenus interact with each other, the FRET can happen under excitation. (**B**) The BiFC-FRET assay of interactions among Pol1-Vc155, Spt16-Vn173 and H3-mCerulean. The fluorescent signals of yeast cells expressing single mCerulean or mVenus, fused mCerulean-mVenus (positive control) and co-expressing mCerulean with mVenus (negative control) were acquired in mCerulean, mVenus, FRET and bright field (BF) channel, respectively. The corrected FRET channel was calibrated as described in Methods. The fluorescence of cells co-expressing H3-mCerluean, Spt16-Vn173 and Pol1-Vc155 were also acquired in all channels. All colors were arbitrarily assigned to indicate signal strength. Scale bar: 3 μm. (**C**) FRET ratio (FR) values obtained from individual yeast cells expressing the indicated fluorescence proteins in (**B**) were calculated. The mean and standard deviation (SD) of FR were shown, with *P* values derived from the Student’s t-test (*** *P*<0.001). (**D**-**G**) BiFC results of Spt16-Pol1 (**D**), Spt16-H3 (**E**), Pol1-H3 (**F**) and Pol1-H2B (**G**) in WT and *spt16-m* cells and the statistical analysis of their BiFC fluorescence signals. The percentages of cells with BiFC fluorescence signals were shown at the lower-left corners of the images. The fluorescence intensity was quantified by ImageJ (number=120). The mean and standard deviation (SD) were shown, with *P* values derived from a nonparametric t-test (*** *P*<0.001). Scale bar: 3 μm. A.U., arbitrary units. (**H**) Pull-down of Pol α with FACT-yH3-H4 or FACT-yH2A-H2B *in vitro*. Pol α was incubated with pre-bound FACT-yH3-H4 or FACT-yH2A-H2B complex. The proteins bound with FACT complex were detected by Western blotting using specific antibodies. yH3-H4 or yH2A-H2B were added with the indicated amounts and two titrations of Pol α were incubated in each concentration of histones. Different amounts of purified proteins were loaded as input groups.

We further combined BiFC with the fluorescence resonance energy transfer (FRET) (Fig. 1A). Spt16 and Pol1 were fused with Vn173 and Vc155, respectively. When FACT and Pol α interact with each other, BiFC emission is generated under 514 nm excitation. H3 was fused with mCerulean. If H3 interacts with the FACT-Pol α complex, excitation at 440 nm will trigger FRET between mCerulean and mVenus, generating BiFC emission (Fig. 1A). The corrected FRET signal was calculated to reflect the BiFC-FRET efficiency. Expressing mCerulean or mVenus alone (Fig. 1B, Panels 1 and 2), or co-expressing them (Fig. 1B, Negative control, Panel 3) did not trigger corrected FRET signals (Fig. 1B and 1C). Fusion of mVenus and mCerulean served as a positive control (Fig. 1B, Panel 4, and 1C). Corrected FRET signals were detected in the S phase with Spt16::Pol1-mVenus and H3-mCerulean (Fig. 1B, Panel 5, and 1C), demonstrating that Spt16, H3-H4, and Pol1 interact with each other in S phase. These signals were significantly higher in S phase cells than in non-S phase cells (Fig. 1B, compare Panel 5 and Panel 6, and 1C), indicating a temporal and spatial specificity of the Pol α-H3-H4-FACT ternary complex formation.

FACT is the histone chaperone for both H3-H4 and H2A-H2B (*22, 42*). Pol1 also binds both H3-H4 and H2A-H2B (*39, 41*). To determine the specific role of histones in the FACT-Pol α interaction, we tested the interactions in the *spt16-m* background (*spt16-K692AR693A* in the Spt16 middle domain), which showed a significant defect in the binding of FACT with H3-H4, but not H2A-H2B (*43*). Notably, BiFC signals of Spt16::Pol1, Spt16::H3, and Pol1:: H3 were significantly reduced in *spt16-m* mutant cells (Fig. 1 D-F and S1C-E). However, we did not detect an apparent reduction of BiFC signals in the Pol1::H2B pair in *spt16-m* mutant cells (Fig. 1G and S1F). These results indicate that histone H3-H4 is crucial for the formation of the FACT-Pol α and Pol α-H3-H4 complexes.

To further test this idea, we performed *in vitro* pull-down assays using purified yeast H3-H4, H2A-H2B, FACT complex and Pol α complex (fig. S2 and Fig. 1H). FACT was first incubated with either H3-H4 or H2A-H2B, followed by the addition of Pol α. The GFP-associated protein complex was then eluted and analyzed by Western blot. Both H2A-H2B and H3-H4 were detected in the Spt16-GFP-associated complex (Fig. 1H). Importantly, Pol1-CBP signals were detected only when FACT was pre-incubated with H3-H4 (Fig. 1E, compares lanes 12-15 with lanes 6-11). With equal amounts of FACT, higher Pol1-CBP signals were observed with increasing amounts of H3-H4 (Fig. 1H, compares lanes 12-13 with lanes 14-15). These results collectively demonstrate that H3-H4, but not H2A-H2B, is essential for forming the FACT-histones-Pol α ternary complex.

### The FACT-(H3-H4)-Pol α ternary complex stimulates Pol α activity

To assess whether FACT-(H3-H4)-Pol1 interaction regulates Pol1’s enzymatic activity, we performed a primer extension assay (Fig. 2A). A 70 nt poly-dT template annealed with a Cy3-labelled 15 nt poly-dA primer was used as the DNA substrate (Fig. 2B). Pol α and FACT complexes were purified from yeast (Fig. 2C). After adding the Pol α complex, DNA products were resolved on a denaturing PAGE gel at each time point (Fig. S3A and Fig. 2D). Products ranging from ∼15 (primer length) to 70 nt (full length products) were generated, with a sharp band at ∼70 nt increasing in intensity over time (fig. 3, A and B, and Fig. 2, D-F). Notably, adding the preformed FACT-(H3-H4) complex significantly increased the intermediate products and the sharp band at 70 nt (Fig. 2, D-F), indicating that the FACT-(H3-H4) complex significantly stimulates Pol1 activity. When using the FACT complex with the *spt16-m* mutation pre-incubated with histone H3-H4, there was a reduction in intermediate products and the full-length product compared to the wild-type FACT complex (Fig. 2, G and H, and fig. S4). These results demonstrate that the FACT complex carrying H3-H4 stimulates the DNA polymerase activity of Pol1.

**Fig. 2.**
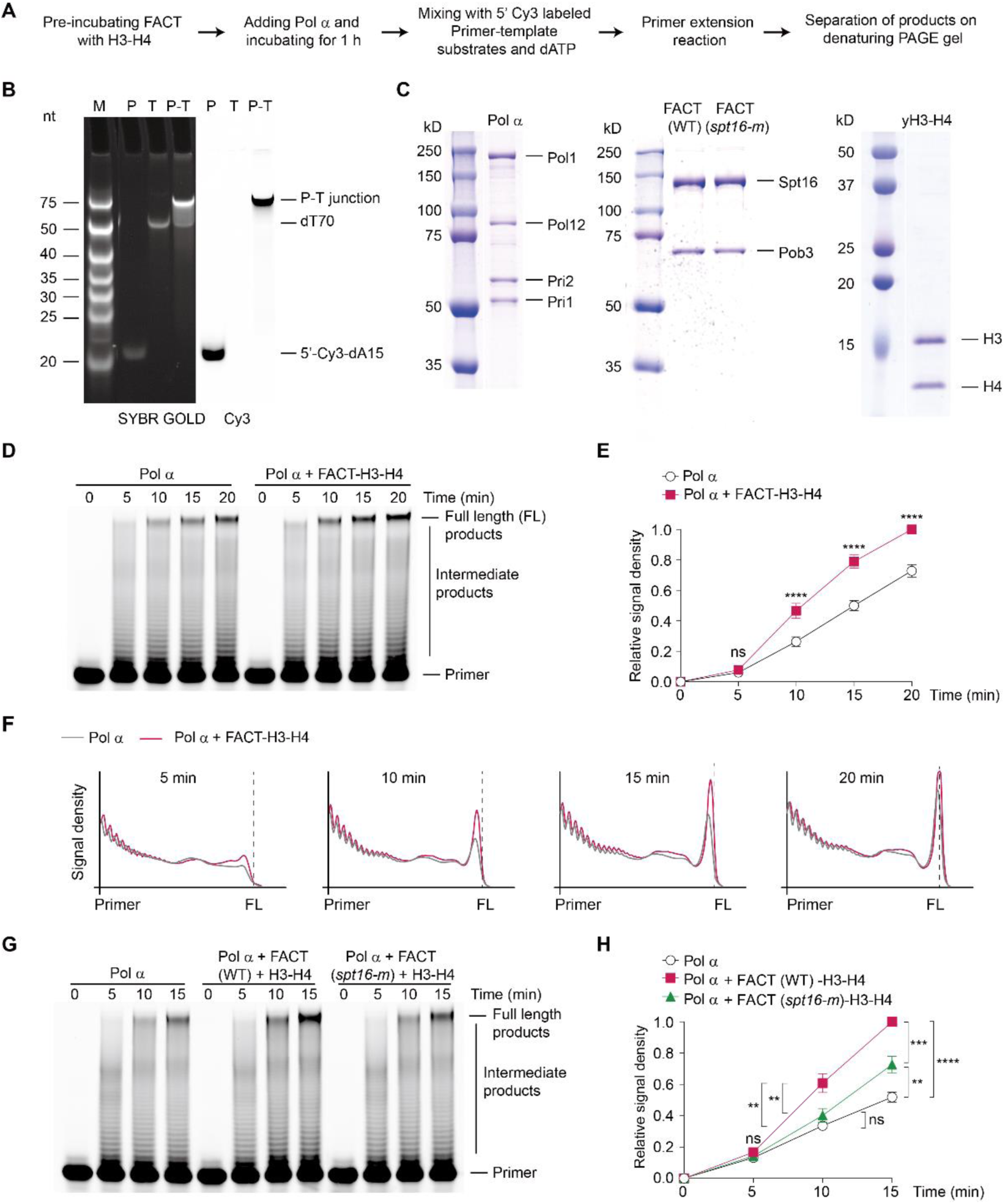
FACT-(H3-H4) promotes primer extension activity of Pol α *in vitro*. (**A**) The scheme of *in vitro* primer extension assay of Pol α. (**B**) Equal moles of 5’-Cy3-labeled dA15 primer and dT70 template were annealed to form the primer-template junction. 5’-Cy3-labeled dA15 primer, dT70 template and annealed primer-template junction are detected on a native gel. Total DNA was stained using SYBR^®^ GOLD, and the Cy3 signals were detected by 568 nm laser. M, DNA marker; P, primer; T, template; P-T junction, primer-template junction. (**C**) Purified Pol α complex, WT or *spt16-m* of FACT complex from yeast cells and yeast H3-H4 purified from *E. coli*, were stained by Coomassie Brilliant Blue (CBB). (**D**) The effect of the FACT-H3-H4 complex on the primer extension activity of Pol α. 5’-Cy3-labeled primer annealed with a template were used as substrates of this assay. The products of this assay at indicated time points were separated on denatured PAGE gel and exposed to a 568 nm laser. The patterns of products are shown in the figure. (**E**) Quantification of the greyscale intensity of full-length products in (**B**). The mean values and standard error (SE) were shown, and the difference between the two groups was analyzed by one-way ANOVA (**** *P*<0.0001, ns *P*>0.05). (**F**) Quantification of the greyscale intensity of products from P (Primer) to FL (Full length products) on the gel in (**D**) at indicated time points. (**G** and **H**) The effect of adding WT or *spt16-m* of FACT with H3-H4 on the primer extension activity of Pol α (**G**) and quantification of full-length products greyscale intensity of it (**H**). The mean values and standard error (SE) were shown, and the difference between the two groups was analyzed by one-way ANOVA (**** *P*<0.0001, *** *P*<0.001, ** *P*<0.01, ns *P*>0.05).

**Fig. 3.**
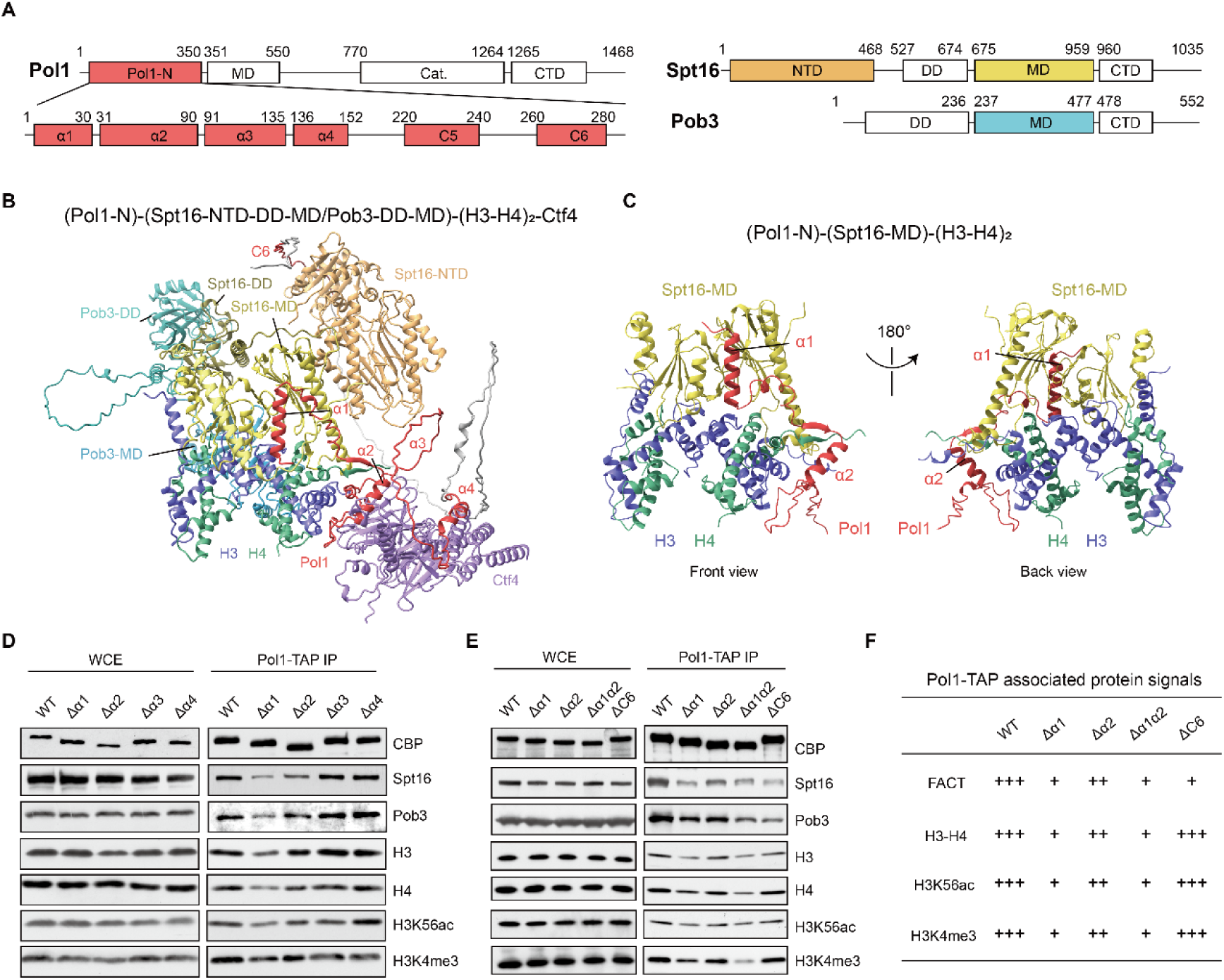
Pol1-N domain is required for the formation of FACT-(H3-H4)-Pol α complex. (**A**) The structural domain of the full length Pol1 subunit in *S. cerevisiae*, the structural domains of the Pol1-N domain predicted by AlphaFold-Multimer 2 and the structural domains of FACT subunits Spt16 and Pob3 in *S. cerevisiae*. (**B**) The predicted structure of *S. cerevisiae* (Pol1-N)-(Spt16-NTD-DD-MD/Pob3-DD-MD)-(H3-H4)_2_-(Ctf4-C) by AlphaFold-Multimer 2. Pol1-N domain is colored in grey, and the four α-helix α1-α4 and coil 6 (C6) of Pol1 are colored in red. Spt16-NTD is colored in orange, Spt16-DD and Spt16-MD are colored in deep and light yellow, respectively. Pob3-DD and Pob3-MD are colored in light blue. H3 and H4 are colored in blue and green, respectively, and Ctf4-C is colored in purple. (**C**) The front (left) and back (right) views of the predicted structure of *S. cerevisiae* (Pol1-N)-(Spt16-MD)-(H3-H4)_2_. Only α1 and α2 of Pol1-N are shown and colored in red. Spt16-MD is colored in yellow, H3 and H4 are colored in blue and green, respectively. (**D**) Pol1-TAP IP to determine the contribution of α1-α4 of Pol1-N domain on the interaction between Pol1 and FACT/(H3-H4)_2_. The proteins in whole cell extracts (WCE) and proteins associated with Pol1-CBP were detected by Western blotting using specific antibodies as indicated. (**E**) Pol1-TAP IP to determine the contribution of α1, α2 and C6 of Pol1-N domain on the interaction between Pol1 and FACT/(H3-H4)_2_. The proteins in whole cell extracts (WCE) and proteins associated with Pol1-CBP were detected by Western blotting using specific antibodies as indicated. (**F**) A table to show the degree of Pol1-FACT/(H3-H4)_2_ interaction displayed in (**D**) and (**E**) in different mutants of Pol1-N domain. +++, strong interaction, ++, middle interaction, +, weak interaction.

### The N-terminus of Pol1 contributes to the FACT-(H3-H4)-Pol1 interaction

The N-terminus of Pol1 (Pol1-N) contains a histone-binding motif (*39, 41*). AlphaFold-Multimer prediction confirmed the Pol1-N domain’s interaction with histone H3-H4 (fig. S5). Considering FACT binding to H3-H4 and Ctf4 interacting with the Pol1 N-terminus, we predicated the interfaces of the Pol1-N domain (1-330 aa), histone H3-H4 tetramer, FACT (Spt16 1-959 aa and Pob3 1-477 aa), and Ctf4-C domain (461-927 aa) (Fig. 3, A and B, and fig. S6). Results show Pol1-N has four alpha helices (red): α1 interacts with the middle domain of Spt16 (Spt16-MD, yellow), α2 interacts with one H3-H4 dimer (H3-H4, blue and green), and α3 and α4 interact with Ctf4 (purple) (Fig. 3B and fig. S6C). Interestingly, the linker sequence upstream of α2 forms an anti-parallel beta-sheet with the N-terminal tail of H4 (Fig. 3B). Additionally, a random coil domain, C6, interacts with the N-terminus of Spt16 (Spt16-NTD, orange) (Fig. 3B and fig. S6C). Predicted structure of Pol1-N domain, H3-H4 tetramer and Spt16-MD (675-959 aa) also show Pol1-α1 interacting with Spt16-MD and Pol1-α2 interacting with H3-H4 (Fig. 3C and fig. S7).

To confirm the predicted interface, we purified the Pol α complex from yeast cells with TAP-tagged wild-type or truncated Pol1. We found that both FACT subunits (Spt16 and Pob3), H3 and H4, newly synthesized histones marked with H3K56ac, and parental histone marked by H3K4me3, were detected in wild-type Pol1-TAP complexes (Fig. 3D). Consistent with predictions, signals for histones and FACT were significantly reduced in the Pol1 mutant lacking the α1 domain (*Δα1*), and moderately reduced in the mutant lacking α2 (*Δα2*). No significant changes were observed with deletions of α3 or α4 (*Δα3*, *Δα4*) (Fig. 3, D and F). A dramatic reduction in these signals was observed in the mutant lacking both α1 and α2 (*Δα1α2*) (Fig. 3, E and F), indicating that α1 and α2 helices are critical for the FACT-(H3-H4)-Pol1-N complex formation. Deletion of the C6 region (*ΔC6*) reduced FACT binding but did not affect histone signals. These results demonstrate that α1 and α2 regions of Pol1 are crucial for the interaction among FACT, histone H3-H4, and Pol1, while the C6 region may only contribute to FACT binding.

### The FACT-(H3-H4)-Pol α interaction is required to maintain replication speed and Okazaki fragment synthesis

To further investigate the role of FACT-(H3-H4)-Pol1 interaction in DNA replication, we then assessed cell cycle progression and DNA replication speed in mutant cells with disrupted FACT-(H3-H4)-Pol1 interactions. Cells lacking α1 and α2 motifs in Pol1-N (*pol1*-*Δα1α2*) showed a significant delay of S-phase progression (Fig. 4A). Using DNA combing assay, we monitored DNA replication fork progression and origin selection (Fig. 4B). Cells were arrested at G1 phase, released into S phase, and nascent DNA was labeled with incorporated BrdU. Each BrdU track represents two sister forks emerging from a single origin, and the BrdU track length reflects the average fork speed (Fig. 4B). Our results showed a significant, but with different degrees of reduction in the average length of BrdU tracks in *Δα1* (∼16.60 kb), *Δα2* (∼14.96 kb) and *Δα1Δα2* (∼14.37 kb) mutant cells compared to wild-type cells (∼19.28 kb), but not in *ΔC6* (∼18.24 kb) cells, indicating the decreased replication fork progression in Pol1 mutants lacking interaction with H3-H4 (Fig. 4C).

**Fig. 4.**
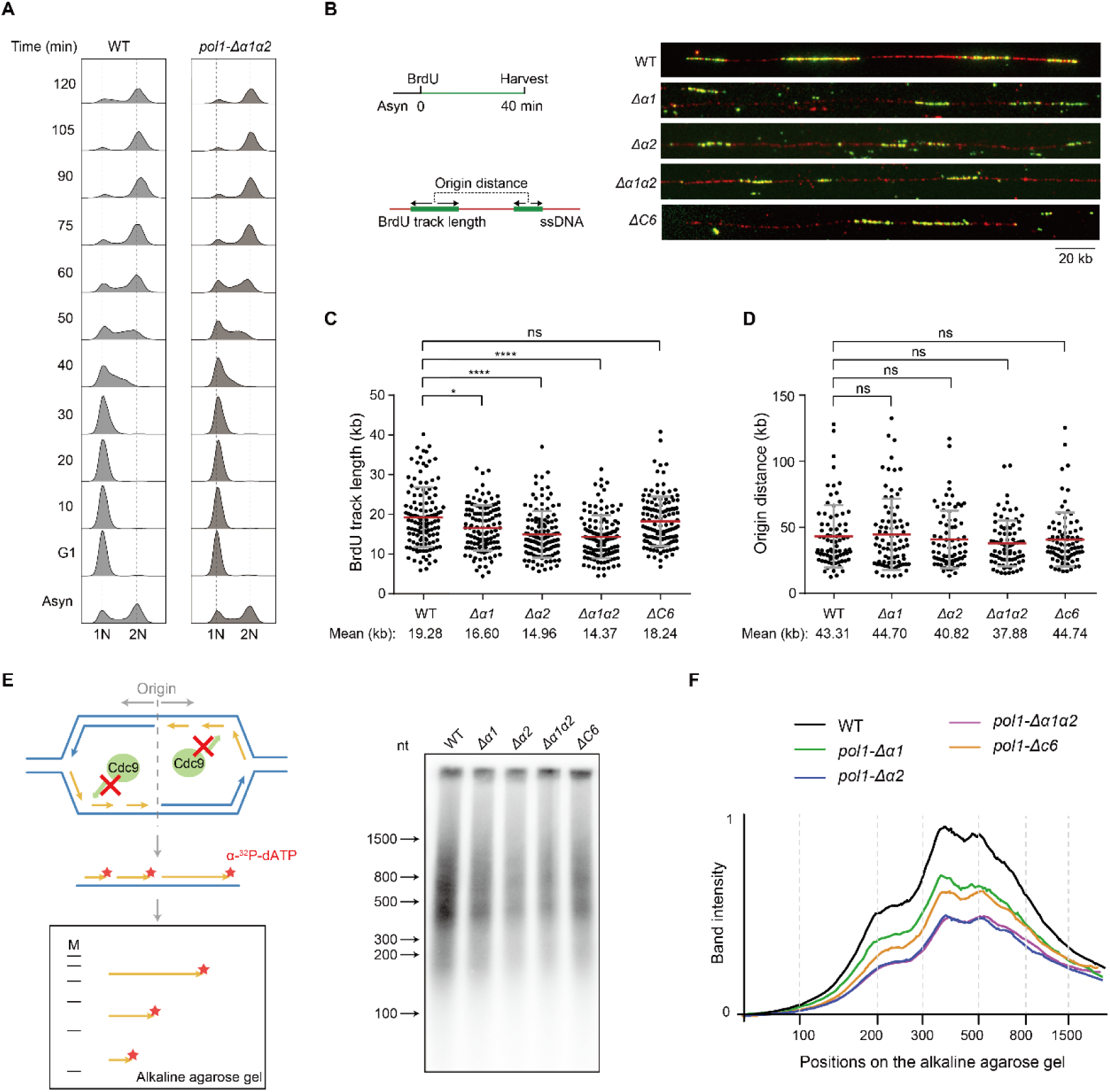
The FACT-(H3-H4)_2_ binding ability of Pol1-N domain is required for replication elongation and Okazaki fragment synthesis. (**A**) Cell cycle progression of wild-type (WT) and *pol1-Δα1α2* cells. Cells were collected at indicated time points after release from G1 phase, and DNA contents were then monitored by flow cytometry. (**B**) The scheme of DNA combing assay (left) and representative DNA tracks of WT, *pol1-Δα1*, *pol1-Δα2*, *pol1-Δα1α2* and *pol1-ΔC6* cells (right). Left: Schematic of DNA combing assay to monitor replication fork progression in S phase with BrdU labeling. Each BrdU tract indicates two divergent replication forks that start from the same origin. Center-to-center distances of BrdU tracks indicate the average replication origin distances. Representative images of BrdU tracks (green) and full-length DNA (red) in WT and Pol1-N mutant cells are shown (right). (**C** and **D**) Dot plots of BrdU track lengths (**C**) and distances of BrdU track centers (**D**) in WT, *pol1-Δα1*, *pol1-Δα2*, *pol1-Δα1α2* and *pol1-ΔC6* cells. The mean and standard deviation (SD) are shown in the figures, and the mean values are indicated at the bottom of the panel. The significance test of differences between the groups was performed using a nonparametric t-test (**** *P*<0.0001, * *P*<0.05, ns *P*>0.05). (**E**) The scheme of Okazaki fragment purification (left) and the result of Okazaki fragment synthesis in WT, *pol1-Δα1*, *pol1-Δα2*, *pol1-Δα1α2* and *pol1-ΔC6* cells by autoradiography (right). Left: In Cdc9-degraded cells, the Okazaki fragments on the lagging strand cannot be ligated efficiently. After purification of the genomic DNA containing un-ligated Okazaki fragments, α-^32^P-dATP is used to label the end of each DNA fragment, and the DNA separated on alkaline gel can be detected by autoradiography. Right: The purified Okazaki fragments of WT and Pol1-N domain mutant cells, the marker of DNA size is labeled at the left of the panel. (**F**) The grey scale intensity of each lane in (**e**) measured by ImageJ with the size of DNA marker.

To rule out the reduced replication fork progression due to altered origin firing, we measured the density of fired origins by analyzing the average distance between origin centers (Fig. 4D). Our analysis showed no significant difference in origin distance between WT and Pol1 mutants, indicating that these Pol1 mutations do not affect replication origin selection (Fig. 4D).

Pol α initiates Okazaki fragment synthesis for lagging strand replication. We then purified Okazaki fragments from the above Pol1-N mutant cells with DNA ligase I Cdc9 degradation (*44*). The un-ligated Okazaki fragments were end-labeled with α-^32^P-dATP, separated on denaturing agarose gel, and detected by autoradiography (Fig. 4E). Purified Okazaki fragments were significantly reduced in *Δα1*, *Δα2,* and *Δα1Δα2* mutant cells, and moderately reduced in *ΔC6* mutant cells compared to wild-type cells (Fig. 4E and F). Together, these results demonstrate that the FACT-(H3-H4)-Pol1 interaction is critical for maintaining replication fork progression and Okazaki fragment synthesis in cells.

### The FACT-(H3-H4)-Pol α interaction senses replication-coupled nucleosome assembly level to adjust the replication speed

To further assess the role of FACT-(H3-H4)-Pol α interaction in the regulation of replication, we assessed the replication rate in *spt16-m* mutant cells. We found that *spt16-m* mutant cells exhibited a delay in S phase progression (fig. S8A). We then used the DNA combing assay to monitor DNA replication fork progression and origin selection under unperturbed conditions (fig. S8B). Our results showed a significant reduction in the average length of BrdU tracks in *spt16-m* mutant cells (∼13.77 kb) compared to wild-type cells (∼19.31 kb) (fig. S8C), indicating a decrease in replication fork progression in *spt16-m* relative to wild-type. No significant difference in the average distance of origins between WT and *spt16-m* cells (∼40 kb) was detected (fig. S8D), suggesting that *spt16-m* mutation does not affect the overall selection of replication origin. Moreover, the *spt16-m* mutant cells showed a significant reduction in purified Okazaki fragments (fig. S9, A and B). Thus, the ability of FACT to bind H3-H4 is critical for the formation of the FACT-(H3-H4)-Pol α complex, which is necessary for Okazaki fragment synthesis to maintain an appropriate DNA replication fork progression rate.

Besides FACT, the histone H3-H4 chaperones CAF-1 and Rtt106 are also involved in replication-coupled nucleosome assembly at replication fork regions (*45–47*). To test whether these chaperones can interact with Pol1, TAP-tagged Cac2 (a subunit of CAF-1), Rtt106, and Spt16 (a subunit of FACT) were purified from yeast cells expressing Flag-tagged Pol1 (Fig. 5A). H3K56ac signals were detected in all three chaperone complexes, Pol1-Flag signals were only detected in the Spt16-TAP complex and not in the Cac2 or Rtt106-TAP complexes (Fig. 5A). Consistently, no BiFC signals were observed in Pol1::Cac1 or Pol1::Rtt106 pairs (Fig. 5B). Of note, cell cycle progression delay is also observed in *cac1*Δ and/or *rtt106*Δ mutant cells (*18, 46*). To determine whether the FACT-(H3-H4)-Pol1 interaction is affected in these histone chaperone mutant cells, we purified the Pol1-TAP complex from cells lacking CAF-1 and/or Rtt106. Importantly, both FACT and histone signals in the Pol1-associated complex were significantly reduced in *cac1*Δ and *cac1*Δ *rtt106*Δ mutant cells, and were mildly reduced in *rtt106Δ* mutant cells. We also measured BiFC signals of Pol1::H3 in *cac1*Δ and/or *rtt106*Δ mutant cells (Fig. 5D). The results showed that Pol1-H3 interaction was disturbed in *cac1*Δ and *rtt106*Δ, especially the double mutant cells (Fig. 5D and E). Thus, we believe that the FACT-(H3-H4)-Pol α interaction may provide a general mechanism for sensing multiple histone chaperone-mediated replication-coupled nucleosome assembly levels to maintain the efficient replication fork progression.

**Fig. 5.**
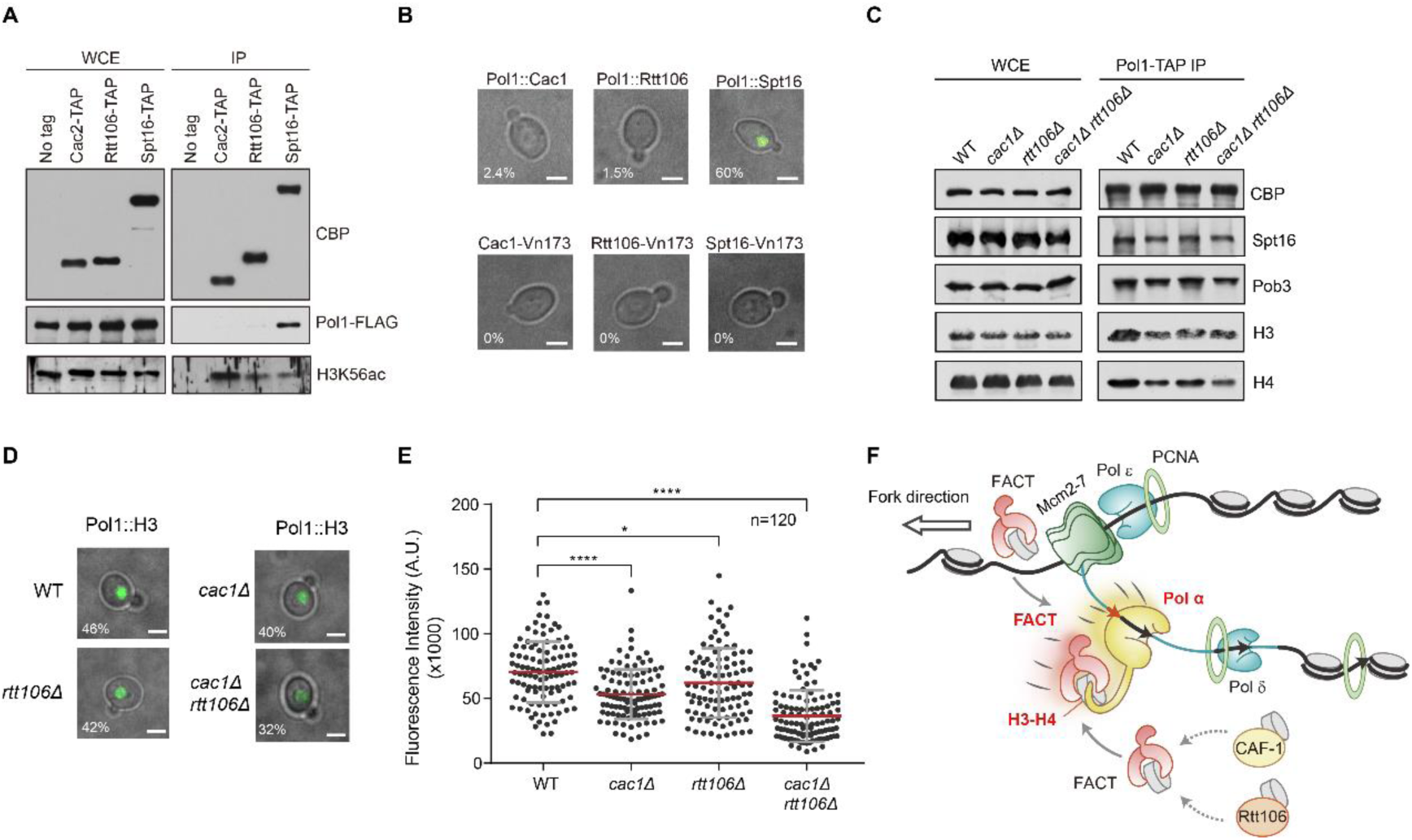
CAF-1 and Rtt106 regulate the formation of FACT-(H3-H4)-Pol α complex. (**A**) Using Cac2-TAP, Rtt106-TAP and Spt16-TAP IP to purify Pol α. The proteins in whole cell extracts and those associated with each histone chaperones were detected using specific antibodies. WCE, whole cell extracts. (**B**) BiFC results of Cac1-Pol1, Rtt106-Pol1 and Spt16-Pol1 interactions in yeast cells. Pol1 was fused with Vc155, while Cac1, Rtt106 and Spt16 were fused with Vn173. The expression of all fusion proteins was under their intrinsic promoters. The cells only carrying Cac1, Rtt106 or Spt16-Vn173 were used as the negative control. The percentages of cells with BiFC fluorescence signals were shown at the lower-left corners of the images. Scale bar: 3 μm. (**C**) Pol1-TAP IP to purify histone H3-H4 and FACT complex in WT, *cac1Δ*, *rtt106Δ* and *cac1Δ rtt106Δ* strains. The proteins in whole cell extracts and proteins associated with Pol1-CBP were detected with specific antibodies. WCE, whole cell extracts. (**D**) BiFC results of Pol1-H3 interaction in WT, *cac1Δ*, *rtt106Δ* and *cac1Δ rtt106Δ* strains. The percentages of cells with BiFC fluorescence signals were shown at the lower-left corners of the images. Scale bar: 3 μm. (**E**) The statistical analysis of BiFC signal intensity in (**D**). The fluorescence intensity in cells was quantified by ImageJ (number=120). The mean and standard deviation (SD) were shown, with *P* values derived from a nonparametric t-test (**** *P*<0.0001, * *P*<0.05). A.U., arbitrary units. (**F**) Model of FACT-(H3-H4)-Pol α function as Pre-Warning System. Histone H3-H4 mediates the interaction between FACT and Pol α, and the formation of FACT-(H3-H4)-Pol α complex is critical for stimulating Pol α activity to maintain Okazaki fragment synthesis and DNA replication speed. CAF-1 and Rtt106-mediated RC nucleosome assembly can also affect the FACT-(H3-H4)-Pol α complex formation and regulate DNA replication. As the FACT-(H3-H4)-Pol α complex formation monitors synchronization between nucleosome assembly and DNA replication, thus we refer to this mechanism as a Pre-Warning System (PWS).

## Discussion

Our research has uncovered a novel mechanism for regulating DNA replication through the FACT-(H3-H4)-Pol α interaction. We demonstrated that histone H3-H4, but not H2A-H2B, is required for the interaction between FACT and Pol α during the S phase. Moreover, the FACT-(H3-H4) complex stimulates the DNA polymerase activity of Pol α to maintain replication fork progression. Disruption of this interaction results in reduced Okazaki fragment synthesis and slowed replication progression speed. Remarkably, this interaction is also regulated by CAF-1 and/or Rtt106-mediated RC nucleosome assembly pathways. Together, these results indicate that the FACT-(H3-H4)-Pol α interaction serves as a general mechanism to orchestrate DNA synthesis with RC nucleosome assembly pathways at the replication fork (Fig. 5F).

Nucleosome assembly is a stepwise process, in general, wherein the initial step involves histone H3-H4 tetramer binding with DNA, followed by the subsequent addition of two H2A-H2B dimers. We demonstrated that histone H3-H4, but not H2A-H2B, mediates the interaction between FACT and Pol α complex. FACT consists of multiple domains and can interact with both histone H3-H4 and H2A-H2B. We and others have demonstrated that the Spt16-MD is required for the interaction between FACT and histone H3-H4 (*23, 43, 48, 49*). Considering that H3-H4 assembly is the first step for nucleosome assembly, an insufficient supply of H3-H4 means that DNA replication-coupled nucleosome assembly would be defective. If the DNA replication speed is not properly adjusted, newly synthesized DNA would remain unprotected by nucleosomes, leading to severe DNA damage. Given H2A-H2B assembly occurs in subsequent steps and H2A-H2B is highly dynamic in cells, it is not a suitable mediator for reflecting nucleosome assembly status compared to H3-H4. Thus, the involvement of histone H3-H4 in promoting the FACT-Pol α interaction further reinforced the intrinsic link of regulating the enzymatic activity of Pol α to synchronize with histone availability. When a sufficient amount of histone H3-H4 carried by FACT is bound to Pol α, it facilitates the synthesis of primers, promoting Okazaki fragment DNA synthesis and DNA replication fork progression.

A previous study has shown that histone H3-H4 promotes the formation of Asf1-(H3-H4)-Mcm2-7 complex ahead of the replication fork, and provides a mechanism to regulate parental DNA unwind activity by histone supply (*12, 50*). While it was observed that DNA replication is impaired in Asf1 knockdown cells, it remained unclear how Asf1-(H3-H4) binding impacts replisome helicase activity. Here, we showed that histone H3-H4 bridged FACT-(H3-H4)-Pol α interaction regulates Pol α activity in response to chromatin assembly pathways. More importantly, this interaction serves as a general mechanism in response to multiple histone chaperones, specifically all H3-H4 chaperones directly connected to the replication fork (FACT, CAF-1, and Rtt106), mediating replication-coupled chromatin assembly pathways. Viewed in combination, these studies demonstrated that multiple pathways in DNA replication can be monitored to fine-tune the DNA synthesis in response to the nucleosome assembly status. This precautionary measure is crucial for preventing severe DNA damage resulting from the exposure of DNA, therefore, we refer to it as a Pre-Warning System (PWS) (Fig. 5F).

We believe the PWS system senses the DNA replication-coupled histone supply to monitor the nucleosome assembly status. This system fine-tunes the DNA replication speed to ensure synchronization between DNA synthesis and nucleosome assembly. Activation of the PWS prevents the situation where newly synthesized naked DNA loses the protection of nucleosomes, and emphasizes that avoiding a potentially dire situation is as crucial as the DNA repair system in safeguarding our genetic material and preserving genome integrity.

## Acknowledgments

We thank all members of the Li lab for technical and intellectual support throughout the project.

## Funding

This work was supported by the National Natural Science Foundation of China (NSFC 31830048, 31725015 to Q.L., NSFC 32370623 to J.F.), and the Beijing Outstanding Young Scientist Program (BJJWZYJH01201910001005 to Q.L).

## Author Contributions

Conceptualization: W.Z., Q.L., J.F;

Methodology: W.Z., J.X., J.Y., J.W., Q.L., J.F;

Investigation: W.Z., J.X., J.Y., G.S., N.G., Q.L., J.F;

Funding acquisition: Q.L., J.F;

Project administration: Q.L., J.F;

Supervision: Q.L., J.F;

Writing – original draft: W.Z., J.F., Q.L.

## Competing interests

The authors declare that they have no competing interests.

## Data and materials availability

All strains and plasmids are available upon request.

## Supplementary Materials

Materials and Methods

Figs. S1 to S9

Tables S1 to S3

References

## Notes

### Competing Interest Statement

The authors have declared no competing interest.

