## Supplementary Information for "FACT-(H3-H4) complex stimulates Pol α activity to coordinate DNA synthesis with nucleosome assembly"

The Supplementary Materials includes:

Materials and Methods

Figs. S1 to S9

Tables S1 to S3

References

### Materials and Methods

#### Yeast strains and Culture conditions

All *S. cerevisiae* yeast strains used in this study were derived from W303-1 *(leu2-3,112 trp1-1 can1-100 ura3-1 ade2-1 his3-11,15*). All genotypes, plasmid constructions, and oligos were listed in Supplementary Tables 1-3. YPD and SC (Synthetic Complete) media were used to culture all yeast strains.

#### DNA combing assay

The method was modified from previous studies (*51*, *52*). Yeast cells were cultured in YPD media at 30°C to $OD_{600}$ about 0.5, arrested at the G1 phase by α-factor (5 mg/mL, 1000X) at 25 °C. Cells were spun down at 2,500 rpm for 5 min at 4°C, washed twice with cold $ddH_2O$ and then released into fresh YPD media with 0.4 mg/mL BrdU at 23°C for 40 minutes to label nascent DNA. Cells (about 50 $OD_{600}$) were collected, washed twice with cold $ddH_2O$, and washed once with cold buffer z (1.2 M sorbitol, 50 mM Tris-HCl pH 7.4). The pellets were resuspended with 4.35 mL buffer z (10 mM β-mercaptoethanol freshly added) and digested with 106.5 μL zymolyase at 28°C for about 35 min to get spheroplasts. The efficiency of digestion was evaluated by measuring $OD_{600}$ in 1% SDS, with a decrease of over 90%. The pellets were then resuspended with 500 μL buffer z (pre-warmed at 42°C) and mixed with 500 μL 1% LMP (low melting point) agarose gel (melted at 65°C and kept at 42°C). 90 μL of cell plugs were generated in casting molds (90 μL) and solidified at 4°C for 30 min. The plugs were then digested in proteinase K buffer (10 mM Tris–HCl, pH 7.5, 50 mM EDTA, 1% sarkosyl, 2 mg/mL proteinase K) twice for 2 days (2 mL proteinase K buffer for 5 plugs) at 42°C. After the digestion, plugs were washed with $TE_{50}$ (10 mM Tris-HCl, pH 7.0, 50 mM EDTA, pH 8.0) 5 times for 10 min and can be stored at 4°C in $TE_{50}$ for a week. One plug was picked up each time and washed once with 5 mL 1×MES buffer, pH 6.2 (50 mM MES) for 5 min. 1×MES buffer was diluted from 500 mM MES, pH 6.2, which was adjusted by adding 500 mM MES sodium to 500 mM MES hydrate. The plug was then transferred to 300 μL fresh 1×MES buffer, pH 6.2, and melted at 65°C for 15 min. After cooling down, 3 μL of β-agarase I was added to the solution, and the digestion was at 42°C for overnight. The solution was spun down at 13,300 rpm on the following day. 40 μL of DNA solution in 1×MES buffer, pH 6.2 was added onto a clean glass slide and a silanized coverslip (Genomic Vision, COV-001) was covered in the solution and incubated for 5 min. Then the DNA was stretched at a speed of 300 μm/s and cross-linked at 60°C for 2 h. DNA was fixed in 70%, 90%, and 100% ethanol for 5 min respectively, then denatured in 1 M NaOH for 22 min and neutralized with 1×PBS. BrdU was detected with a rat monoclonal antibody followed by a goat anti-rat coupled to Alexa 488, and ssDNA was detected using anti-ssDNA and goat anti-mouse coupled to Alexa 546. Images of DNA fibers were recorded with DragonFly (Andor). The number of DNA fibers calculated per sample was over 120, two biological replicates were scored and statistically analyzed, and one representative result was shown.

**Yeast dot assay**

To analyze phenotypes of yeast strains, serial dilution (5 or 10-fold) of fresh cultures concentrated to $OD_{600}$ of about 0.6 were spotted onto YPD plates containing different concentrations of drugs. Plates were inoculated at 30°C or other temperatures for the time noted in each figure legend. Each dot assay was performed with three replicates and one representative result was shown.

**Trichloroacetic Acid (TCA) precipitation for whole cell extract to detect Rad53 phosphorylation**

Yeast cells were collected and treated with TCA lysis buffer (1.85 M NaOH, 7.4% β-mercaptoethanol) for 10 min on ice, then followed by adding 20% TCA and inverting, with another 10 min on ice. The sample was centrifuged and the supernatant was removed. The pellets were washed with acetone (pre-cooled at -20°C) and dried in a concentrator. Then, the sample was dissolved in 0.1 M NaOH buffer and an equal volume of SDS sample buffer. The whole cell extract sample was resolved by SDS-PAGE gel and the signals were detected by Western blotting with indicated antibodies.

**Tandem Affinity Purification (TAP)**

TAP-tagged purifications were performed as described (*53, 54*). Briefly, exponentially growing yeast cells were harvested by centrifugation and washed with 10% glycerol. Yeast cells were resuspended in equal volumes of buffer A (25 mM Tris, pH 8.0, 100 mM NaCl, 1 mM EDTA, 10 mM $MgCl_2$, 0.01% NP40) containing protease inhibitors (1 mM DTT, 1 mM PMSF, 1 mM Benzamidine, 1 mM Pefabloc) and 15 kU/ml DNase I, and frozen in liquid nitrogen. Frozen cells were ground in a freezer mill (SPEX SamplePrep$^{LLC}$ 6870). Cell lysates were clarified by centrifugation (20,450 g, 40 minutes) and then incubated with IgG sepharose beads (GE Healthcare) for 2 hours at 4°C. IgG-bound proteins were digested with TEV protease for 2 hours at 16°C after extensive washing. The resulting proteins were further incubated with calmodulin beads (Agilent Technologies), eluted with SDS sample buffer, resolved on SDS-PAGE gel, and detected by Western blotting.

**Protein purification for *in vitro* assays**

yFACT complex purification: GFP tag was integrated into the C-terminal of the *SPT16* gene at its endogenous genomic locus. The GFP purification was performed as previously described with minor modifications (*24*). Briefly, yeast cells were resuspended in equal volumes of buffer A with 200 mM NaCl and treated 15 KU/mL DNase I. Cell lysates were incubated with GFP nanobody beads for 2 hours at 4°C. After binding, the beads were washed 3 times with buffer A with 500 mM NaCl to remove nonspecifically associated proteins. The beads coupled with the FACT complex were stored at 4°C.

Yeast DNA polymerase α complex purification for *in vitro* pull-down: TAP tag was integrated into the C-terminal of the *POL1* gene at its genomic locus. The TAP purification was performed as described above with minor modifications. Briefly, yeast cells were resuspended in equal volumes of buffer A with 100 mM NaCl and treated with 15 KU/mL DNase I. Cell lysates were incubated with IgG beads for 2 hours at 4°C and the beads were washed 3 times with buffer A with 300 mM NaCl. The IgG beads bound proteins were digested with TEV protease, and the eluted DNA polymerase α complex solution was stored at 4°C.

yH3-H4 purification: BL21 (DE3) strain carrying yH3-H4 expression vector was grown to reach $OD_{600}$ to 0.6-0.8. IPTG was added to a final concentration of 0.4 mM, with induction at 37°C for 1 h. Cells were washed once with 1×PBS, resuspended in buffer C (10 mM Hepes, pH7.6, 1 mM EDTA, 10% glycerol, 1 mM DTT, 0.1 mM PMSF, 2 mM Benzamide, 0.1 M NaCl) and sonicated for 3 times, 30 s each time, to break cells. The solution was spun down at 10,000 rpm for 10 min at 4°C. The pellets were resuspended in 0.25 M HCl using a homogenizer and incubated at 30°C for 20 min. The solution was then spun down at 10,000 rpm for 10 min. The supernatant was collected and neutralized with about 0.125 volume of 2 M Tris Base. The solution was dialyzed in buffer C overnight at 4°C and then loaded onto a Source 15S column. The peak was collected at about 1 M NaCl concentration. The purified proteins were stored at -80°C.

yH2A-H2B purification: BL21 (DE3) strain carrying yH2A expression vector and JM109 (DE3) were grown in LB media at 37°C to reach $OD_{600}$ about 0.6-0.8. IPTG was added to a final concentration of 0.4 mM. H2A was induced at 37°C for 1 h and H2B was induced at 37°C for 16 h. Cells were washed with cold 1×PBS and spun down at 30,000 rpm for 15 min at 4°C. The supernatant was collected and neutralized with a 2 M Tris base. The solution was dialyzed overnight in buffer C at 4°C and then loaded onto a Source 15S column. The peak was at about 1 M NaCl concentration and the eluted proteins were stored in -80°C.

Purification of Pol α from yeast cells for primer extension assay: The protocol was followed as previously described (*55*). Briefly, cells were grown to $OD_{600}$ at about 2.0, and expression of Pol α was induced at 30°C for 2 h by the addition of 2% galactose. Cells were collected, washed, and resuspended in buffer D (25 mM Tris-HCl pH7.2, 10% glycerol, 0.02% NP40-S, 1mM DTT) with 400 mM NaCl and protease inhibitors (0.3 mM PMSF, 7.5 mM benzamidine, 0.5 mM AEBSF, 1 mM leupeptin, 1 mM pepstatin A and 1 mg/ml aprotinin). The frozen cells were ground in a freezer mill and the resulting powders were then thawed in buffer D with 400 mM NaCl and protease inhibitors. The mixture was spun down at 235,000 g, for 1 h at 4°C. The concentration of NaCl was reduced to 300 mM by dilution in a buffer lacking NaCl. The solution was then incubated with 2 mM calcium chloride and calmodulin affinity resin at 4°C for 90 min. Then the beads were washed with buffer D with 300 mM NaCl and 2 mM $CaCl_2$, and proteins were eluted with buffer D with 300 mM NaCl, 2 mM EDTA, and 2 mM EGTA. Eluted fractions were diluted in buffer D to 120 mM NaCl and were loaded onto a 5 mL MonoQ column. Proteins bound were eluted and dialyzed in buffer D with 150 mM NaCl.

***In vitro* pull-down assay**

For GFP pull-down, the FACT complex coupled GFP nanobody beads were washed 3 times with buffer A200 (25 mM Tris-HCl pH 7.5, 200 mM NaCl, 1 mM

EDTA, and 0.01% Triton X-100) and incubated with indicated proteins in buffer A200 for 2 h at 4°C. After extensive washing, the beads were incubated with another indicated protein in buffer A200 overnight at 4°C. Protein-bound beads were eluted with SDS sample buffer, resolved by SDS-PAGE gels, and detected by Western blotting.

**Chromatin Immunoprecipitation-quantitative PCR assay (ChIP-qPCR)**

Briefly, exponentially growing cells (*MATa*) were synchronized with 5 μg/ml α-factor for 2 h at 25°C and then released into YPD media with or without 200 mM HU (Sigma). Samples were collected at indicated time points and temperatures for the following ChIP assay. The procedure of following ChIP was described previously (*56*) with minor modifications based on special requirements. Cells were incubated with 1% formaldehyde (Sigma) at 25°C for 20 min and quenched with 0.125 M Glycine at 25°C for 5 min. After beads-beating with glass beads added, the protoplasts were sonicated by a Biorupter (Diagenode) to chromosomal DNA fragments about 200-500 bps in length, and then immunoprecipitated with specific antibodies in demand. All ChIP DNA was enriched by applying indicated antibodies except TAP-tagged ChIP requires IgG beads (GE Healthcare) without further recognition by Protein G Sepharose (GE Healthcare). After extensive washing, the DNA sample was boiled with 10% Chelex-100 (Bio-rad) for reverse crosslinking. The final ChIP DNA products were analyzed by Real-time PCR (CFX96; BioRad) with primers amplifying replication origins. The percentage of ChIP DNA relative to total input DNA was calculated.

**Bimolecular Fluorescence Complementation (BiFC)**

Vn173 and Vc155 fragments were inserted into genes of interest, that were linked with GGGGSGGGGS, respectively. For Spt16-H3 BiFC, Vn173 was integrated into the C-terminal of Spt16 with its endogenous promoter on a plasmid, and Vc155 was integrated into the C-terminal of H3 with $P_{GAL1}$ promoter on plasmid. For Spt16-Pol1 BiFC, Spt16-Vn173 was constructed as described above, and Vc155 was integrated into the C-terminal of Pol1 with $P_{GAL1}$ promoter on the plasmid. For Pol1-H3 BiFC, Vn173 was integrated into the C-terminal of H3 at its genomic loci, and Pol1-Vc155 was constructed as described above. The BiFC assays in (Fig. 5B) was performed by fusing Vc155 to Pol1 and Vn173 to Cac1 subunit of CAF-1, Rtt106 and Spt16 subunit of FACT. Pol1-Vc155 expression was under the intrinsic promoter of $P_{POL1}$ and the expression of Cac1-Vn173, Rtt106-Vn173 and Spt16-Vn173 were under $P_{CAC1}$, $P_{RTT106}$ and $P_{SPT16}$. The yeast strain carrying *POL1-VC155* was crossed with yeast strains carrying fusion Vn173 strains respectively. For constitutive expression, yeast cells were cultured in SC media with 5% glucose, then collected for imaging. For induced expression, yeast cells were cultured in SC media with 2% raffinose and 0.02% glucose. Cells were then transferred into SC media with 2% Galactose, with 3 h of induction at 25°C, followed by imaging. All the data is acquired on Ultraview Vox Spinning Disk Confocal Microscopy (Perkin Elmer) and DragonFly (Andor).

**Intensity-based FRET measurement with three filter microscopies**

Yeast cells containing the BiFC-FRET system were collected and plated onto 0.17 mm thick dishes with a glass slide at the bottom. Then images were taken with an Olympus IX81 inverted microscope equipped with a 100×, NA=1.45 oil immersion objective lens and cool-coupled device. Excitation light was delivered by an X-cite light source. For image acquisition, the Metamorph software was used. In all experiments, the excitation intensity was attenuated down to 25% of the maximal power of the light source. Images were acquired using 1×1 binning mode and 400 ms integration times. For quantitative FRET measurements, the method of sensitized FRET was previously described in detail (*57-59*). Images were acquired sequentially through YFP, FRET, and CFP filter channels. Here, the filter sets were YFP channel (Excitation filter: 504/12 nm; Dichroic mirror: 400/521/607/700 nm; Emission filter: 529/39 nm; Semrock), FRET channel (Excitation filter: 427/10 nm; Dichroic mirror: 400/521/607/700 nm; Emission filter: 529/39 nm; Semrock), CFP channel (Excitation filter: 427/10 nm; Dichroic mirror: 400/521/607/700 nm; Emission filter: 472/30 nm; Semrock). The background images were subtracted from the raw images before carrying out the FRET calculation. Corrected FRET ($F^C$) was calculated on a pixel-by-pixel basis for the entire image using the following equation: $F^C$ =FRET-($a$×YFP)-($b$×CFP), where YFP, FRET and CFP correspond to background subtracted images of cells co-expressing YFP and CFP acquired through the YFP, FRET and CFP channels, respectively. The "*a*" and "*b*" are the fractions of bleed-through of YFP and CFP fluorescence through the FRET filter channel, respectively. To quantify FRET efficiency, we used FR=[FRET-($b$×CFP)]/($a$×YFP), a relative value that varies with changes in energy transfer to quantify the FRET signal. The FRET ratio (FR) represents the fractional increase in YFP emission due to FRET. Thus, in the absence of energy transfer, FR has a predicted value of 1 (*60*).

***In vitro* primer extension assay of Pol α**

The assay was performed as previously described with some modifications (*31*). Briefly, a 15 nt DNA primer, dA15, was annealed with a 70 nt DNA template, dT70, in TE buffer (pH 8.0). The annealed primer-template junction was resolved on 20% native PAGE gel for quality control. To perform the primer extension assay, the reaction was assembled in the following buffer: 50 mM Tris (pH 8.0), 0.05 mg/mL BSA, 5 mM $MgCl_2$, and 1 mM DTT. Purified FACT and yH3-H4 proteins were first incubated in the reaction buffer at 4°C for 1 h, and then Pol α was added into the buffer. The reaction started at 37°C by adding 1 mM annealed primer-template junction and 0.1 mM dATP. The reaction was quenched by adding 2.5 volume of denaturing loading buffer (90% formamide, 10 mM EDTA, and 5 mg/mL bromophenol blue), heated at 70°C for 10 min, and then snap-cooled on ice to keep single-stranded DNA. The sample was loaded onto 18% denaturing urea PAGE gel, exposed upon 568 nm, and stained with SYBR® GOLD. The grey value of all products separated on the gel was measured by ImageJ.

**Budding yeast Okazaki fragment purification and *in vitro* labeling**

Cells carrying Cdc9-AID* degron were cultured to $OD_{600}$ about 0.4. IAA (3-Indoleacetic acid) was added to the final concentration of 1 mM and induced at 30°C for 3 h. 50 mL cells were collected, spun down at 3000 g for 5 min and washed with 10 mL SCE buffer (1 M sorbitol, 100 mM sodium citrate, 60 mM EDTA, pH 7.0). 5 mg 20T zymolase together with β-mercaptoethanol was added to digest the cell wall at 28°C for 30 min. Then, the spheroblasts were washed with 10 mL SCE buffer and digested by adding 480 µl lysis buffer with 150 µg proteinase K (Fisher) at 37°C for 2-16 h. The remaining proteins or peptides were precipitated by adding 200 µl 5.0 M KOAc and spun down at 16000 g for 30 min. The supernatant was collected and added 500 µl isopropanol to precipitate the DNA, spun down at 16000 g for 10 min. The pellet DNA was washed twice with 500 µl 70% ethanol and spin down at 16000 g for 1min. Then, the pellet was resuspended using 200 µl STE buffer and 25 µg RNase A was added to digest RNA at 37°C for 1 h. The DNA was precipitated by adding 20 µl 3 M NaOAc (pH 5.5) and 800 µl ethanol, then was spun down at 5000 g for 10 min. The pellet was washed once with 1 mL ethanol and finally resuspended using 50 µl TE buffer. The DNA sample should be stored at 4°C and cannot be frozen.

For end labeling of the Okazaki fragment by radioactive probes, the total reaction system is 20 µl, containing 5 U Klenow (exo$^-$) polymerase, 33 nM α-$^{32}$P-dATP and 2 µl DNA sample. The reaction is at 37°C for 30 min and then, inactivated at 75°C for 20 min. Using Illustra micro spin G-50 columns (GE healthcare) to remove the free radioactive probes and elute the DNA sample. Alkaline loading buffer was added to the DNA sample and 10 µl sample was loaded to 2% alkaline agarose gel to separate DNA at 150 V for 3 h (Using ice to keep the buffer cool). After electrophoresis, DNA was transferred to the positively-charged nitrocellulose film (Hybond-N$^+$; GE healthcare) overnight. The film was exposed to the phosphor screen and scanned by Typhoon FLA9500 imager (GE healthcare). Using ImageJ to quantify the grey scale intensity of bands.

**Statistics and reproducibility**

At least two independent experiments were performed to generate each dataset. Statistical analysis was performed by Graphpad prism 9. The statistical test used for each experiment is stated in the figure legends.

Fig. S1

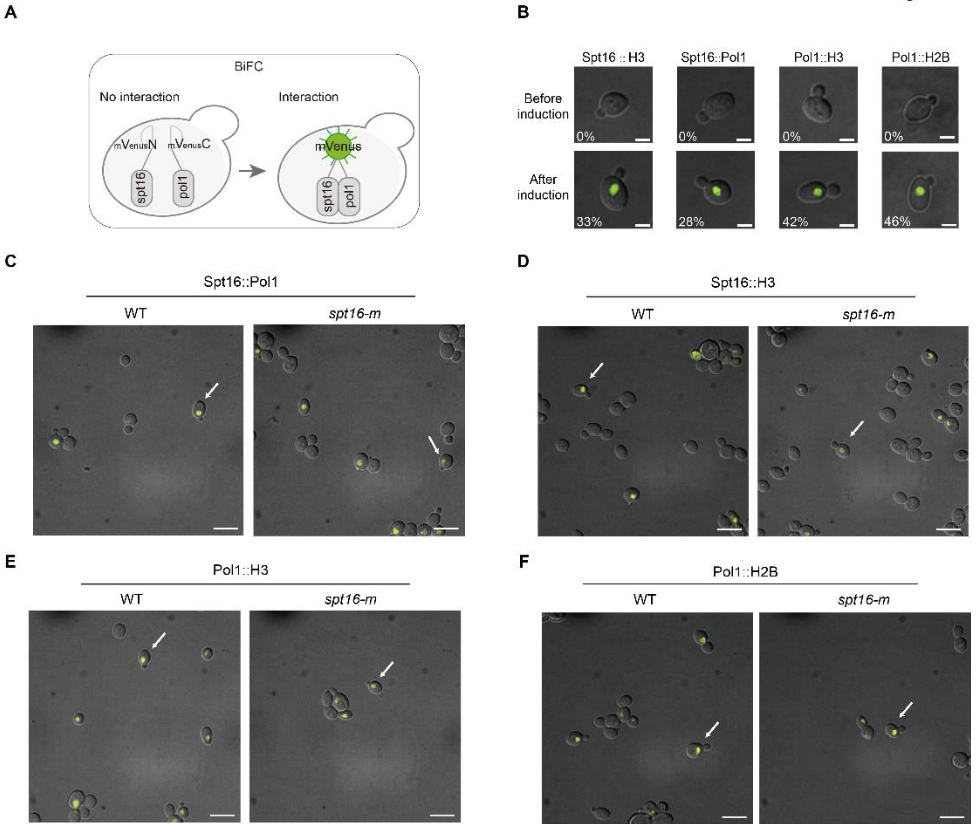

**Fig. S1. Using the BiFC method to visualize interactions of Spt16-H3, Spt16-Pol1, Pol1-H3 and Pol1-H2B in cells.**

(**A**) Schematic of BiFC (Bimolecular fluorescence complementation) method. N- and C-terminal of YFP mVenus were fused to two proteins of interest respectively. Vn173 and Vc155 would reconstruct a functional mVenus when two proteins interact with each other.

(**B**) The BiFC assay to show the interactions of Spt16-H3, Spt16-Pol1, Pol1-H3 and Pol1-H2B in WT cells. The BiFC signals were detected after induction of the corresponding proteins described in the **Materials and Methods**. The images of cells before induction were used as negative control groups. The percentages of cells with BiFC fluorescence signals were shown at the lower-left corners of the images. Scale bar: 3 μm.

(**C-F**) The BiFC of Spt16-H3, Spt16-Pol1, Pol1-H3 and Pol1-H2B in WT and *spt16-m* cells. The representative cells shown in **Fig. 1** were indicated by white arrows. Scale bar: 5 μm.

Fig. S2

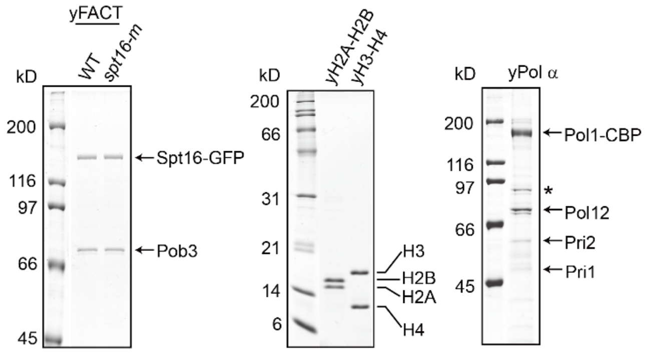

**Fig. S2. The purified proteins used in pull-down assay.**

The GFP and TAP tag were used to purify FACT complex and Pol α complex respectively in yeast. yH2A-H2B and yH3-H4 were purified from *E. coli*. The quality of purified proteins was confirmed by Coomassie Brilliant Blue (CBB) staining. The asterisk (*) indicates the non-specific band.

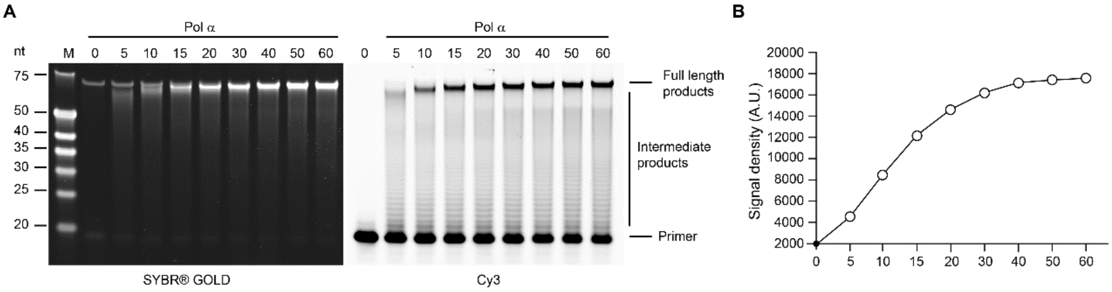

**Fig. S3. The purified Pol α complex displays primer extension activity *in vitro.***

(**A**) The purified Pol α complex catalyzes primer extension *in vitro*. The products of primer extension assay in the presence of purified Pol α at indicated time points were separated by denaturing urea PAGE gel. The total DNA were stained by SYBR® GOLD (Left panel), and the newly generated DNA products were detected by 568 nm laser (Right panel).

(**B**) Quantification of the grayscale intensity in Cy3 channel of the full-length product bands in (**A**) using ImageJ.

Fig. S4

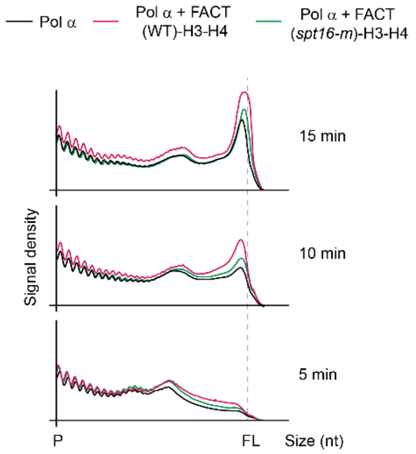

**Fig. S4. *spt16-m* attenuates the promotion of Pol α primer extension activity by FACT-H3-H4 *in vitro.***

The effect of WT and *spt16-m* of FACT complex together with H3-H4 on the primer extension activity of Pol α. The Quantification of grey value of products ranging from P (Primer) to FL (Full Length products) at indicated time points in **Fig. 2D** was shown.

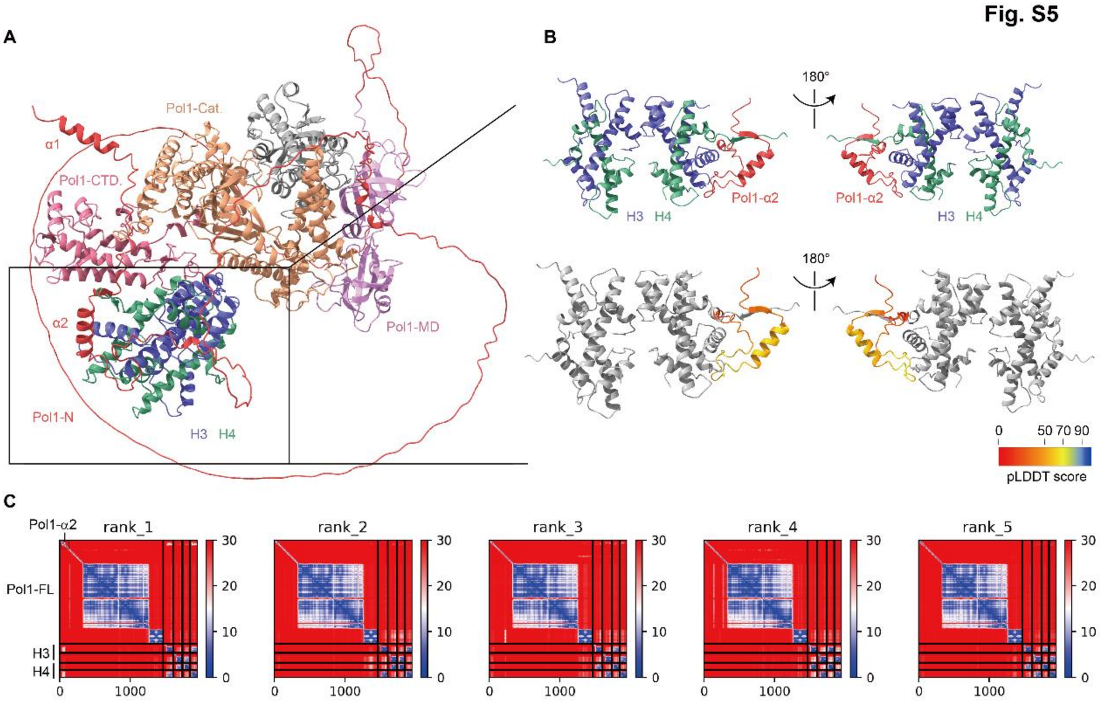

**Fig. S5. AlphaFold-Multimer v2 predicted structures of full-length *S. cerevisiae* Pol1 with histone H3-H4 tetramer.**

(**A**) The first rank of the predicted structure of Pol1-(H3-H4)$_2$ indicates the $\alpha$2 helix of Pol1 N-terminus domain (amino acids 31-90) is predicted to bind to one side of (H3-H4)$_2$.

(**B**) The predicted binding interface between Pol1-N domain with (H3-H4)$_2$ that is magnified from the black square part in (**A**), in which the interaction interface is similar with the one predicted using Pol1-N domain only with (H3-H4)$_2$. Top: The Pol1-$\alpha$2-(H3-H4)$_2$ interaction surface. Pol1-$\alpha$2 is colored in red, H3 and H4 are colored in blue and green, respectively. Bottom: The predicted local-distance difference test (pLDDT) score of the predicted Pol1-$\alpha$2 structure in the top panel, H3 and H4 are colored in grey.

(**C**) PAE plots of all five predicted structures of Pol1-(H3-H4)$_2$ suggests Pol1-$\alpha$2 specifically interacts with (H3-H4)$_2$.

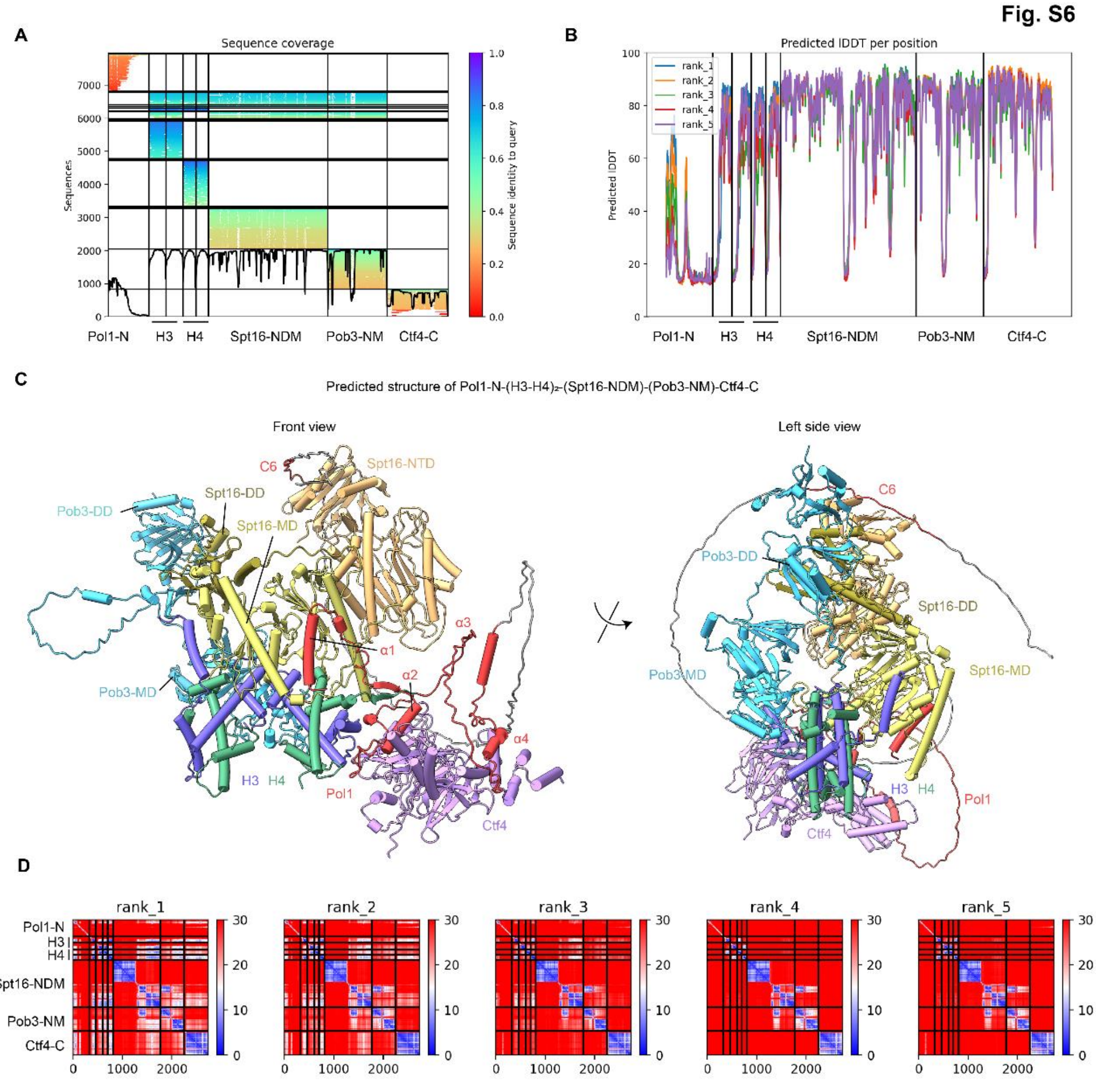

**Fig. S6. The predicted structures of *S. cerevisiae* (Pol1-N)-(Spt16-NTD-DD-MD/Pob3-DD-MD)-(H3-H4)$_2$-(Ctf4-C) by AlphaFold-Multimer v2.**

(**A**) AlphaFold-generated multiple sequence alignment results of *S. cerevisiae* Pol1-N domain, histone H3 and H4, Spt16-NTD-DD-MD, Pob3-DD-MD and Ctf4-C domain between species. The color of each alignment indicates the similarity between the input sequence and aligned sequence (scale from 0 to 1).

(**B**) AlphaFold pLDDT score of all amino acids of Pol1-N domain, histone H3 and H4, Spt16-NTD-DD-MD, Pob3-DD-MD and Ctf4-C domain. High pLDDT score indicates that confidence for the location of the amino acid is high.

(**C**) The front (left) and side (right) view of the predicted (Pol1-N)-(Spt16-NTD-DD-MD/Pob3-DD-MD)-(H3-H4)$_2$-(Ctf4-C) structure. Pol1-N domain is colored in grey and the four α-helix α1-α4 and coil 6 (C6) of Pol1 is colored in red. Spt16-NTD is colored in orange, Spt16-DD and Spt16-MD are colored in deep and light yellow, respectively. Pob3-DD and Pob3-MD are colored in light blue. H3 and H4 are colored in blue and green, respectively, and Ctf4-C is colored in purple.

(**D**) PAE plots of all five predicted structures of (Pol1-N)-(Spt16-NTD-DD-MD/Pob3-DD-MD)-(H3-H4)$_2$-(Ctf4-C) suggests that Pol1-α1 specifically interacts with Spt16-MD, Pol1-α2 specifically interacts with (H3-H4)$_2$ and Pol1-α4 specifically interacts with Ctf4.

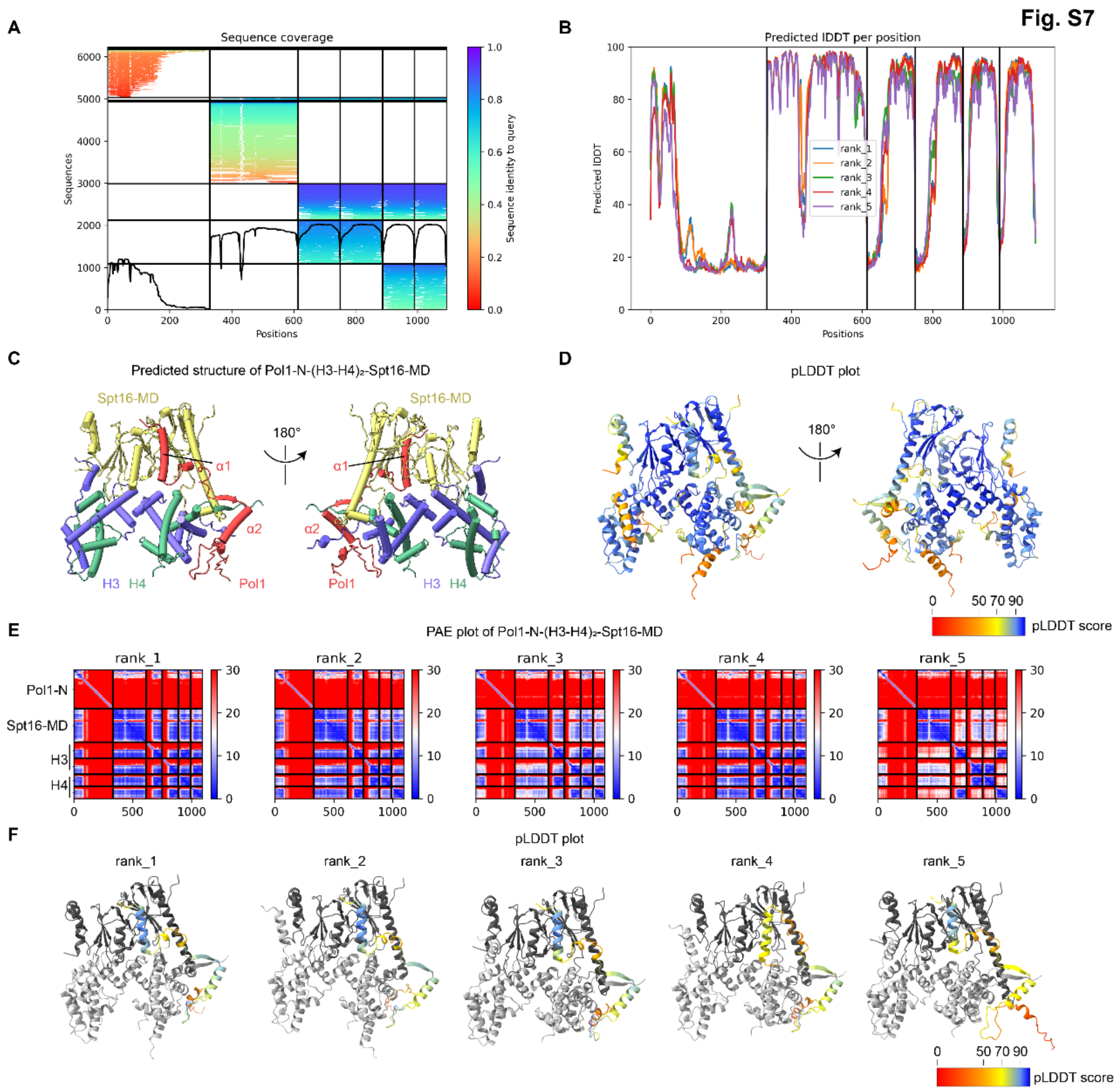

**Fig. S7. Quality control of the predicted structures of *S. cerevisiae* (Pol1-N)-(Spt16-MD)-(H3-H4)$_2$ by AlphaFold-Multimer v2.**

(**A**) AlphaFold-generated multiple sequence alignment results of *S. cerevisiae* Pol1-N domain, histone H3 and H4 and Spt16-MD between species. The color of each alignment indicates the similarity between the input sequence and aligned sequence (scale from 0 to 1).

(**B**) AlphaFold pLDDT score of all amino acids of Pol1-N domain, histone H3 and H4, and Spt16-MD. High pLDDT score indicates that confidence for the location of the amino acid is high.

(**C**) The front (left) and back (right) view of the predicted (Pol1-N)-(Spt16-MD)-(H3-H4)$_2$ structure. Pol1-α1 and α2 are colored in red, Spt16-MD is colored in yellow, H3 and H4 are colored in blue and green, respectively.

(**D**) The pLDDT score of the predicted (Pol1-N)-(Spt16-MD)-(H3-H4)$_2$ structure. High pLDDT score indicates that confidence for the location of the amino acid is high.

(**E**) PAE plots of all five predicted structures of (Pol1-N)-(Spt16-MD)-(H3-H4)$_2$.

(**F**) The pLDDT score of Pol1-α1 and α2 in all five predicted structures of (Pol1-N)-(Spt16-MD)-(H3-H4)$_2$ show that the predicted interaction surface here is similar to that of (Pol1-N)-(Spt16-NTD-DD-MD/Pob3-DD-MD)-(H3-H4)2-(Ctf4-C) (**Fig. S6**).

Fig. S8

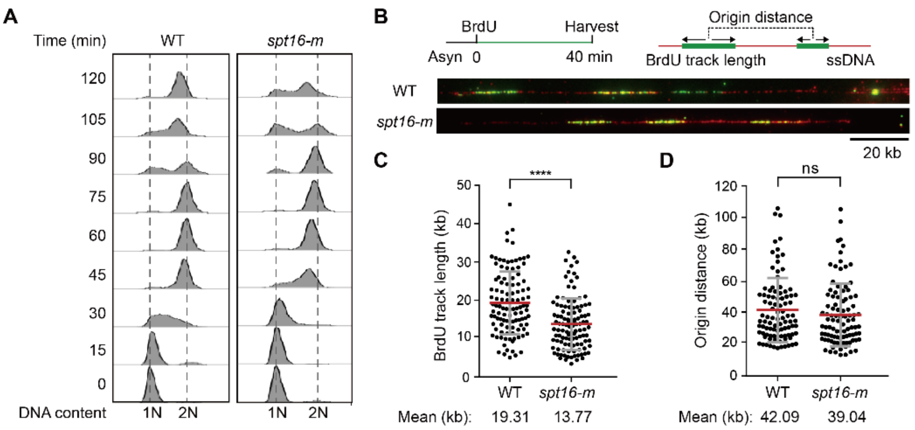

**Fig. S8. *spt16-m* mutation leads to reduced replication elongation rate.**
(**A**) Cell cycle progression of wild-type (WT) and *spt16-m* cells is shown. Cells were collected at indicated time points after release from G1 phase, and DNA contents were then monitored by flow cytometry.
(**B**) The scheme of DNA combing assay and representative DNA tracks of WT and *spt16-m* cells. Top: Schematic of replication fork progression assay with BrdU labeling. Each BrdU tract indicates two divergent replication forks that start from the same origin. Center-to-center distances of BrdU tracks indicate the average replication origin distances. Bottom: Representative images of BrdU tracks (green) and full-length DNA (red) in WT and *spt16-m* cells are shown.
(**B** and **C**) Dot plots of BrdU track lengths (**C**) and distances of BrdU track centers (**D**) in WT and *spt16-m* cells. The mean and standard deviation (SD) are shown in the figure, and the mean values are indicated at the bottom. The significance test of differences between the two groups was performed using a nonparametric t-test (**** $P$<0.0001, ns $P$>0.05).

Fig. S9

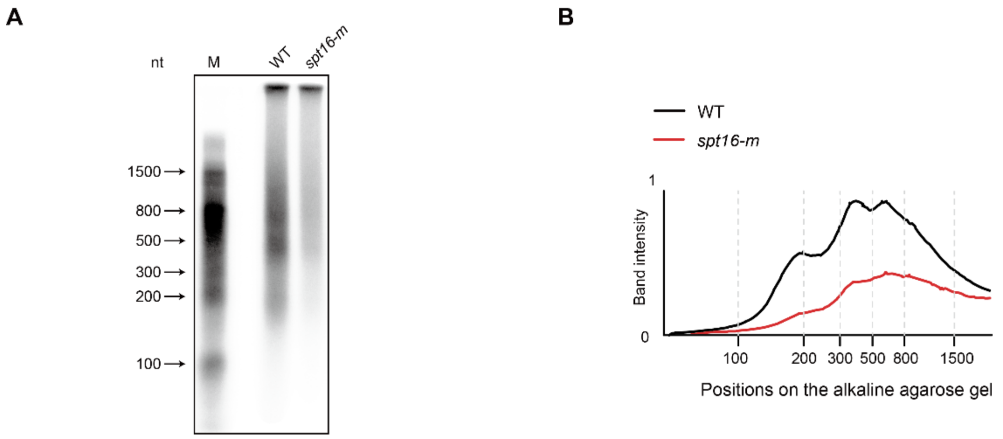

**Fig. S9. *spt16-m* mutation leads to defective Okazaki fragment synthesis.**
(**A**) The purified and α-$^{32}$P-dATP labeled Okazaki fragments of WT and *spt16-m* mutant cells. The purification procedure is the same as in **Fig. 4E**. Purification of the genomic DNA containing un-ligated Okazaki fragments from Cdc9-degraded cells, α-$^{32}$P-dATP is used to label the end of each DNA fragment, and the DNA separated on alkaline gel was detected then by autoradiography. The marker of DNA size is labeled at the left of the panel.
(**B**) The grey scale intensity of each lane in (**A**) is measured by ImageJ. And the position of DNA marker is indicated in dashed grey line.

**Table S1 Yeast strains used in this study.**

| Name | Genotype |
|---|---|
| LQY170 | *MATa leu2-3/112 ura3-1 trp1-1 his3-11/15 ade2-1 can1-100 rad17Δ::kanMX6* |
| LQY665 | *MATa leu2-3/112 ura3-1 trp1-1 his3-11/15 ade2-1 can1-100 p404-BrdU-Inc::TRP1* |
| LQY819 | *MATa leu2-3/112 ura3-1 trp1-1 his3-11/15 ade2-1 can1-100 p404-BrdU-Inc::TRP1 spt16-m::natR* |
| LQY1248 | *MATa leu2-3/112 ura3-1 trp1-1 his3-11/15 ade2-1 can1-100 mcm4-5Flag::natR* |
| LQY1309 | *MATa leu2-3/112 ura3-1 trp1-1 his3-11/15 ade2-1 can1-100 mcm4-5Flag::natR spt16-m::natR* |
| LQY2293 | *MATa leu2-3/112 ura3-1 trp1-1 his3-11/15 ade2-1 can1-100 CDC45-3HA::kanMX6* |
| LQY2487 | *MATa leu2-3/112 ura3-1 trp1-1 his3-11/15 ade2-1 can1-100 spt16-m::natR* |
| LQY2523 | *MATa leu2-3/112 ura3-1 trp1-1 his3-11/15 ade2-1 can1-100 rad24Δ::kanMX6* |
| LQY2740 | *MATa leu2-3/112 ura3-1 trp1-1 his3-11/15 ade2-1 can1-100 spt16-m::natR rad24Δ::kanMX6* |
| LQY2741 | *MATa leu2-3/112 ura3-1 trp1-1 his3-11/15 ade2-1 can1-100 spt16-m::natR rad17Δ::kanMX6* |
| LQY2852 | *MATa leu2-3/112 ura3-1 trp1-1 his3-11/15 ade2-1 can1-100 cdc45-3HA::kanMX6 spt16-m::natR* |
| LQY3160 | *MATa leu2-3/112 ura3-1 trp1-1 his3-11/15 ade2-1 can1-100 POL1-TAP::TRP1* |
| LQY3184 | *MATa leu2-3/112 ura3-1 trp1-1 his3-11/15 ade2-1 can1-100 spt16Δ::TRP1 pRS313-SPT16-VN173-HIS3* |
| LQY3511 | *MATa leu2-3/112 ura3-1 trp1-1 his3-11/15 ade2-1 can1-100 p404-BrdU-Inc::TRP1 spt16-m::natR* |
| LQY3252 | *MATa leu2-3/112 ura3-1 trp1-1 his3-11/15 ade2-1 can1-100 TAP-SLD5::TRP1* |
| LQY3274 | *MATa leu2-3/112 ura3-1 trp1-1 his3-11/15 ade2-1 can1-100 HHT2-VN173::HIS3* |
| LQY3640 | *MATa leu2-3/112 ura3-1 trp1-1 his3-11/15 ade2-1 can1-100 HHT2-mCerulean::TRP1 spt16Δ::TRP1 pRS313-SPT16-VN173* |
| LQY6056 | *MATa leu2-3/112 ura3-1 trp1-1 his3-11/15 ade2-1 can1-100 pol1-Spt16-MD::HIS3 spt16-m::natR* |
| LQY6151 | *MATa leu2-3/112 ura3-1 trp1-1 his3-11/15 ade2-1 can1-100 PRI1-Flag::natR pol1-Spt16-MD::HIS3* |

| *Continued* | |
|---|---|
| LQY6153 | *MATa leu2-3/112 ura3-1 trp1-1 his3-11/15 ade2-1 can1-100 PRI1-Flag::natR pol1-Spt16-MD::HIS3 spt16-m::natR* |
| LQY6196 | *MATa leu2-3/112 ura3-1 trp1-1 his3-11/15 ade2-1 can1-100 trp1::pRS304-POL1-*$P_{GAL1\text{-}10}$*-Pol12-TRP1 ura3::pRS306-CBP-TEV-PRI1-* $P_{GAL1\text{-}10}$*-PRI2-URA3* |
| LQY6206 | *MATa leu2-3/112 ura3-1 trp1-1 his3-11/15 ade2-1 can1-100 pol1-(spt16-m)-MD::HIS3* |
| LQY6207 | *MATa leu2-3/112 ura3-1 trp1-1 his3-11/15 ade2-1 can1-100 pol1-(spt16-m)-MD::HIS3 spt16-m::natR* |
| LQY6236 | *MATa leu2-3/112 ura3-1 trp1-1 his3-11/15 ade2-1 can1-100 TAP-sld5::KanMX6 cdc45-3 × HA::KanMX6 Pol1-5 × FLAG::natR spt16-m::natR* |
| LQY7761 | *MATa leu2-3/112 ura3-1 trp1-1 his3-11/15 ade2-1 can1-100 pol1-Δα1-TAP::TRP1* |
| LQY7764 | *MATa leu2-3/112 ura3-1 trp1-1 his3-11/15 ade2-1 can1-100 pol1-Δα2-TAP::TRP1* |
| LQY7787 | *MATa leu2-3/112 ura3-1 trp1-1 his3-11/15 ade2-1 can1-100 pol1-Δα3-TAP::TRP1* |
| LQY7789 | *MATa leu2-3/112 ura3-1 trp1-1 his3-11/15 ade2-1 can1-100 pol1-Δα4-TAP::TRP1* |
| LQY7796 | *MATa leu2-3/112 ura3-1 trp1-1 his3-11/15 ade2-1 can1-100 pol1-ΔC6-TAP::TRP1* |
| LQY7832 | *MATa leu2-3/112 ura3-1 trp1-1 his3-11/15 ade2-1 can1-100 pol1-Δα1α2* |
| LQY7837 | *MATa leu2-3/112 ura3-1 trp1-1 his3-11/15 ade2-1 can1-100 pol1-ΔC5-TAP::TRP1* |
| LQY7839 | *MATa leu2-3/112 ura3-1 trp1-1 his3-11/15 ade2-1 can1-100 pol1-Δα1α2C6-TAP::TRP1* |
| LQY7858 | *MATa leu2-3/112 ura3-1 trp1-1 his3-11/15 ade2-1 can1-100 pol1-Δα1 CDC9-AID*::HIS3 ura3::*$P_{ADH1}$*-OSTIR-URA3* |
| LQY7859 | *MATa leu2-3/112 ura3-1 trp1-1 his3-11/15 ade2-1 can1-100 pol1-Δα2 CDC9-AID*::HIS3 ura3::* $P_{ADH1}$*-OSTIR-URA3* |
| LQY7860 | *MATa leu2-3/112 ura3-1 trp1-1 his3-11/15 ade2-1 can1-100 pol1-Δα1α2 CDC9-AID*::HIS3 ura3::* $P_{ADH1}$*-OSTIR-URA3* |
| LQY7863 | *MATa leu2-3/112 ura3-1 trp1-1 his3-11/15 ade2-1 can1-100 pol1-Δα1α2C6* |
| LQY8000 | *MATa leu2-3/112 ura3-1 trp1-1 his3-11/15 ade2-1 can1-100 pol1-Δα1 p404-BrdU-Inc::TRP1* |
| LQY8001 | *MATa leu2-3/112 ura3-1 trp1-1 his3-11/15 ade2-1 can1-100 pol1-Δα2 p404-BrdU-Inc::TRP1* |
| LQY8002 | *MATa leu2-3/112 ura3-1 trp1-1 his3-11/15 ade2-1 can1-100 pol1-Δα1α2 p404-BrdU-Inc::TRP1* |

| Continued | |
|---|---|
| LQY8003 | *MATa leu2-3/112 ura3-1 trp1-1 his3-11/15 ade2-1 can1-100 pol1-ΔC6 p404-BrdU-Inc::TRP1* |
| LQY8004 | *MATa leu2-3/112 ura3-1 trp1-1 his3-11/15 ade2-1 can1-100 pol1-Δα1α2C6 p404-BrdU-Inc::TRP1* |
| LQY8063 | *MATa leu2-3/112 ura3-1 trp1-1 his3-11/15 ade2-1 can1-100 TAP-SLD5::kanMX6 pol1-α1α2-Flag::natR* |

**Table S2: Constructs used in this study**

| Name | Sequence |
|---|---|
| LQP958 | *pRS313-$P_{ADH1}$-NLS-mCerulean* |
| LQP959 | *pRS316- $P_{ADH1}$-NLS-mVenus* |
| LQP962 | *p416/$P_{GPD}$-mCerulean-mVenus* |
| LQP991 | *p426/$P_{GAL1}$-pol1-Vc155* |
| LQP1019 | *p426/$P_{GAL1}$-hht2-Vc155* |

**Table S3: Primers used in this study**

| Name | Sequence |
|---|---|
| ARS607 (F) | TGCCGCACGCCAAACATTGC |
| ARS607 (R) | CGGCTCGTGCATTAAGCTTG |
| ARS305 (F) | AGCAAGACCGGCCAGTTTGA |
| ARS305 (R) | GCACTTTGATGAGGCTCTAGCAA |
